# Glacial connectivity and current population fragmentation in sky-islands explain the contemporary distribution of genomic variation in two narrow-endemic montane grasshoppers from a biodiversity hotspot

**DOI:** 10.1101/2021.01.12.426363

**Authors:** Vanina Tonzo, Joaquín Ortego

**Affiliations:** Department of Integrative Ecology, Estación Biológica de Doñana (EBD-CSIC); Avda. Américo Vespucio 26 – 41092; Seville, Spain

**Keywords:** Approximate Bayesian Computation, ddRAD-seq, demographic history, environmental niche modelling, landscape genetics, Pleistocene glaciations

## Abstract

**Aim:** Cold-adapted biotas from mid-latitudes often show small population sizes, harbor low levels of local genetic diversity, and are highly vulnerable to extinction due to ongoing climate warming and the progressive shrink of montane and alpine ecosystems. In this study, we use a suite of analytical approaches to infer the demographic processes that have shaped contemporary patterns of genomic variation in *Omocestus bolivari* and *O. femoralis*, two narrow-endemic and red-listed Iberian grasshoppers forming highly fragmented populations in the sky island archipelago of the Baetic System.

**Location:** Southeastern Iberia.

**Methods:** We quantified genomic variation in the two focal taxa and coupled ecological niche models and a spatiotemporally explicit simulation approach based on coalescent theory to determine the relative statistical support of a suite of competing demographic scenarios representing contemporary population isolation (i.e., a predominant role of genetic drift) *vs*. historical connectivity and post-glacial colonization of sky islands (i.e., pulses of gene flow and genetic drift linked to Pleistocene glacial cycles).

**Results:** Inference of spatial patterns of genetic structure, environmental niche modelling, and statistical evaluation of alternative species-specific demographic models within an Approximate Bayesian Computation framework collectively supported genetic admixture during glacial periods and postglacial colonization of sky islands, rather than long-term population isolation, as the scenario best explaining the current distribution of genomic variation in the two focal taxa. Moreover, our analyses revealed that isolation in sky islands have also led to extraordinary genetic fragmentation and contributed to reduce local levels of genetic diversity.

**Main conclusions:** This study exemplifies the potential of integrating genomic and environmental niche modelling data across biological and spatial replicates to determine whether organisms with similar habitat requirements have experienced concerted/idiosyncratic responses to Quaternary climatic oscillations, which can ultimately help to reach more general conclusions about the vulnerability of mountain biodiversity hotspots to ongoing climate warming.

## 1. INTRODUCTION

Distributional shifts in response to Quaternary climatic oscillations had a dramatic impact on biogeographical patterns of species diversity, abundance and local endemism (Hewitt, 1996; Sandel et al., 2011). These cycles, in particular the high-amplitude climatic changes characterizing the Middle-Late Pleistocene (Jouzel et al., 2007), have also shaped the distribution and spatial patterns of genetic variation in many organisms (Hewitt, 2000). However, multiple studies have documented considerable heterogeneity across regions and taxa in the demographic consequences of Pleistocene glacial cycles (Hewitt, 2000). On the one hand, the impact of Pleistocene glaciations strongly depended on latitude and regional topography, with extinction-recolonization dynamics at higher latitudes (i.e., “southern richness to northern purity” paradigm; Hewitt, 1996) and elevational shifts and more complex processes of population fragmentation and connectivity at lower latitudes such as the tropics or temperate regions (e.g., “refugia within refugia” concept; Gómez & Lund, 2006). On the other hand, the way organisms respond to climate changes strongly depend on species-specific niche requirements and life-history traits, which define favorable/unfavorable climatic periods (glacials or interglacials; Bennett & Provan, 2008) and their capacity to deal with population fragmentation (e.g., micro-habitat preferences, dispersal capacity, etc.; e.g., Massatti & Knowles, 2016; Papadopoulou & Knowles, 2016). These aspects determined the location and extension of Pleistocene refugia, which have played a predominant role on species’ persistence during unfavorable climatic periods and acted as source populations from which species expanded their ranges at the onset of more favorable conditions (Bennett & Provan, 2008). Temperate species currently inhabiting low elevation areas generally restricted their distributions to southern refugia during glacial phases (i.e., glacial refugia) and expanded during interglacial periods whereas cold-adapted species, nowadays presenting fragmented populations at high elevations/latitudes (i.e., interglacial refugia), had much more widespread distributions in glacial stages (Hewitt, 2000).

Mid- and low-latitude alpine and montane taxa represent small-scale replicates of cold-adapted species living at high latitudes (Hewitt, 1996; 2000). Sky islands are a paradigmatic example of interglacial refugia in which alpine and montane organisms currently form highly isolated populations in mountain ranges separated from each other by intervening valleys with unsuitable environmental conditions (e.g., DeChaine & Martin, 2005; Knowles & Massatti, 2017; McCormack et al., 2008; Mouret et al., 2011). Changing climatic conditions cause suitable habitats to expand or shrink along elevational gradients, leading to alternating periods of population isolation and connectivity (DeChaine & Martin, 2005; Knowles, 2000). Periods of isolation on mountain tops are expected to lead to demographic bottlenecks, genetic drift and divergence among populations, while periods of connectivity allow for dispersal and gene flow among formerly isolated populations (DeChaine & Martin, 2005). Species restricted to sky islands often show unique patterns of population genetic structure resulted from climate-induced distributional shifts (Mouret et al., 2011), processes that in some cases have led to lineage diversification and speciation (Knowles, 2000; Knowles & Massatti, 2017). Therefore, montane habitats are often important hotspots of intraspecific genetic diversity, species richness and local endemism, providing an excellent study system for understanding how climatic changes associated with Pleistocene glacial cycles impacted species distributions, demography and genetic diversification (Mairal et al., 2017). Their study takes particular relevance if we consider that species restricted to sky-islands often show small population sizes, harbor reduced levels of local genetic diversity and are highly vulnerable to extinction due to ongoing climate warming and the progressive shrink of alpine ecosystems (Rubidge et al., 2012).

Southeastern Iberia constitutes an important biodiversity hotspot due to the interplay among a vibrating geological history (Meulenkamp & Sissingh, 2003), an extraordinarily complex topography (Braga et al., 2003), and a limited impact of Quaternary glaciations (Hughes & Woodward, 2017). The presence of vast areas free of permanent ice during the coldest stages of the Pleistocene made this region an important glacial refugium for warm-temperate taxa (Gómez & Lunt, 2007; Hewitt, 2011). At the same time, the mountain ranges of the region (Baetic system; > 3,000 m) constitute important interglacial refugia for several cold-adapted organisms that currently persist in severely fragmented populations at high elevation patches of suitable habitat and contribute disproportionally to the extraordinary rates of local endemism of the area (Mota et al., 2002). This offers an ideal biogeographical setting to jointly study the genetic consequences of long-term population fragmentation in sky islands and assess the timescale at which past distributional shifts have shaped contemporary patterns of genetic variation in alpine organisms from mid latitudes.

Here, we use genomic data obtained via restriction site-associated DNA sequencing (ddRAD seq; Peterson et al., 2012) and a suite of analytical approaches to shed light on the demographic processes underlying contemporary patterns of genetic variation in the grasshoppers *Omocestus bolivari Chopard*, 1939 and *O. femoralis* Bolívar, 1908, two narrow-endemic Iberian species inhabiting high elevation areas in the sky island archipelago of the Baetic System (Presa et al., 2016a, b). The two taxa present adjacent but not overlapping distributions, both occupying severely fragmented open habitats of thorny shrub formations (e.g., *Erinacea sp*., *Festuca* sp., *Juniperus* sp.) located at the upper (>1,500 m) thermoclimatic belts of the Mediterranean region. The distribution range of the two species include sky islands of varying size, ranging from large ones in the main massifs of the region (*O. bolivari*: Sierra Nevada and Sierra de Baza-Filabres; *O. femoralis*: Sierra de Cazorla) to tiny patches of suitable habitat located in isolated mountains with maximum elevations close to the lower altitudinal limit of the species (Figure 1). By using these two species of uniform ecological and life-history traits as independent replicates, we exemplify how the integration of genomic and environmental niche modelling (ENM) data within a comparative framework can help to elucidate the responses of species with similar habitat requirements to processes of population fragmentation/connectivity driven by Pleistocene climatic oscillations and understand the temporal scale at which they contributed to shape their contemporary patterns of genetic variation (Knowles & Massatti, 2017). In a first step, (i) we tested whether observed patterns of genomic variation in the two biological replicates reflect the signals of historical population connectivity (i.e., gene flow during glacial periods) and subsequent colonization and population isolation in sky islands (i.e., genetic drift during interglacial periods) or, rather, if post-glacial population fragmentation and genetic drift after the Holocene have entirely monopolized the genetic makeup of contemporary populations. Specifically, we quantified genetic structure and coupled ecological niche models (ENMs) and a spatiotemporally explicit simulation approach based on coalescent theory to mimic complex demographic processes under a suite of scenarios representing historical gene flow and post glacial colonization in sky islands vs. contemporary population isolation (Ray et al., 2010; e.g., Knowles & Massatti, 2017). The expectations under these alternative scenarios were tested and validated against observed genomic data within an Approximate Bayesian Computation (ABC) framework (e.g., He et al., 2013). In a second step, (ii) we analyzed the impact of sky-island size and the spatial location of contemporary populations on their demographic fate and levels of genetic variation. To this end, we tested the hypothesis that genetic diversity is positively associated with the geographic centrality of populations and habitat patch-size and used demographic reconstructions based on genomic data to determine whether changes in effective population size (*N*_*e*_) over time differed between sky-islands of contrasting size and degree of peripheral location (Lira-Noriega & Manthey, 2014).

**FIGURE 1.**
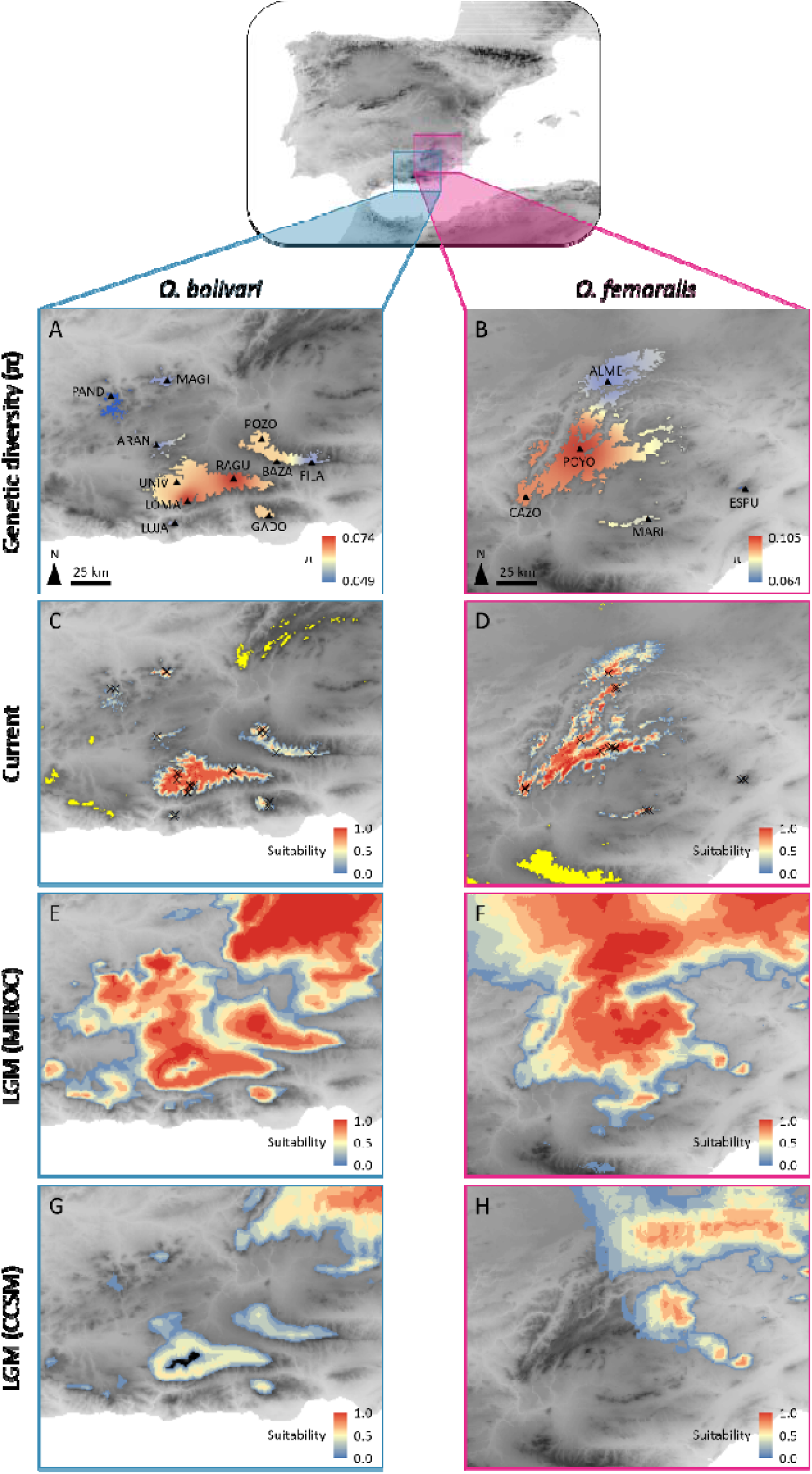
(A-B) Spatial patterns of genetic diversity and (C-H) environmental niche models (ENM) for *Omocestus bolivari* and *O. femoralis*. Panels A-B show genetic diversity (π) of the studied populations (triangles, population codes as in Table S1) interpolated across the current distribution of each species as predicted by their respective ENMs. Panels C-H show the projection of ENMs for (C-D) present and last glacial maximum (LGM) conditions under the (E-F) CCSM4 and (G-H) MIROC-ESM general atmospheric circulation models. Maps for current distributions show records (crosses) used to build the ENMs and dark yellow color indicate areas predicted as suitable out of the known limits of species’ ranges (i.e., over-predictions, not shown in panels A-B). All maps show suitable areas above the maximum training sensitivity plus specificity (MTSS) logistic threshold of MAXENT. Grey background represents elevation, with darker areas corresponding to higher altitudes.

## 2. MATERIALS AND METHODS

### 2.1. Population sampling

Between 2011 and 2016, we sampled eleven populations of *Omocestus bolivari* and five populations of *O. femoralis* (Figure 1; Table S1). According to our extensive surveys in southeastern Iberia and available records in the literature (Presa et al., 2016a, b), the sampled populations cover the entire known distribution range of the two species (Figure 1). We stored specimens in 2 ml vials with 96 % ethanol and preserved them at −20° C until needed for genomic analyses.

### 2.2. Genomic library preparation and processing

We used NucleoSpin Tissue kits (Macherey-Nagel, Düren, Germany) to extract and purify genomic DNA from the hind femur of each individual. We processed genomic DNA into four genomic libraries using the double-digestion restriction-fragment-based procedure (ddRADseq) described in Peterson et al. (2012) (see Tonzo et al., 2019 for details) and used the different programs distributed as part of the STACKS v. 1.35 pipeline (Catchen et al., 2013) to filter and assemble our sequences into *de novo* loci and call genotypes (for details, see Supplementary Methods S1).

### 2.3. Population genetic structure

We analyzed population genetic structure of the two species using the Bayesian Markov chain Monte Carlo clustering method implemented in the program STRUCTURE v. 2.3.3 (Pritchard et al., 2000). We ran STRUCTURE assuming correlated allele frequencies and admixture without using prior population information. We conducted 15 independent runs with 200,000 MCMC cycles, following a burn-in step of 100,000 iterations, for each of the different possible *K* genetic clusters. We retained the ten runs having the highest likelihood for each value of *K* and inferred the number of populations best fitting the dataset using log probabilities [Pr(X|*K*)] (Pritchard et al., 2000) and the Δ*K* method (Evanno et al., 2005). Complementary to STRUCTURE analyses, we performed principal component analyses (PCA) as implemented in the *R* package *adegenet* (Jombart, 2008).

### 2.4. Environmental niche modelling

We built environmental niche models (ENM) to predict the geographic distribution of climatically suitable habitats for *O. bolivari* and *O. femoralis* both in the present and during the last glacial maximum (LGM, 21 ka). Then, we used this information to create maps of environmental suitability in these two time periods and generate alternative demographic models as detailed in the next section (e.g., He et al., 2013; Massatti & Knowles, 2016). To build ENMs, we used the maximum entropy algorithm implemented in MAXENT v.3.3.3 (Phillips et al., 2006) and the 19 bioclimatic variables from the WORLDCLIM dataset (http://www.worldclim.org/) interpolated to 30-arcsec resolution (∼1 km^2^ cell size) (Hijmans et al., 2005). To generate climate suitability maps during the LGM, we projected the ENM onto LGM bioclimatic conditions derived from the CCSM4 (Community Climate System Model; Braconnot et al., 2007) and the MIROC-ESM (Model of Interdisciplinary Research on Climate; Hasumi & Emori, 2004) general atmospheric circulation models. Further details on ENM are presented in Supplementary Methods S2.

### 2.5. Testing alternative demographic models

We used the integrative distributional, demographic and coalescent (iDDC) approach (He et al., 2013) and an Approximate Bayesian Computation (ABC) framework (Beaumont et al., 2002) to test alternative scenarios representing different hypotheses about how landscape heterogeneity (or its lack thereof) and colonization from glacial refugia *vs*. contemporary population isolation in sky islands explain the spatial distribution of genomic variation in our two focal species. This approach is described in He et al. (2013) and consists of three main steps: (i) constructing alternative demographic models representing different hypotheses about the processes shaping spatial patterns of genetic structure and diversity; (ii) running demographic and genetic simulations under each model using the software SPLATCHE2 (see Ray et al., 2010); (iii) evaluating the fit of observed genomic data (i.e., empirical genomic data obtained after genotyping sampled populations) to the genetic expectations under each model, identifying the most probable model/s, and estimating demographic parameters (e.g., He et al., 2013; González-Serna et al., 2019). Below we describe the most relevant aspects of this approach and present an extended version with all the details in Supplementary Methods S3.

#### Constructing demographic models

We generated two main sets of models that differ in the hypothetical demographic processes that have shaped spatial patterns of contemporary genetic variation (Table 1):

**TABLE 1.** Statistics from the ABC procedure used for evaluating the relative support of each demographic model in *Omocestus bolivari* and *O. femoralis*. A higher marginal density correspond to a higher model support and a high Wegmann’s *p*-value indicates that the model is able to generate data in agreement with empirical data. Bayes factors represent the degree of relative support for the model with the highest marginal density over the other models. *R*^2^ is the coefficient of determination from a regression between each demographic parameter (*K*_MAX_, *m, N*_ANC_) and the five partial least squares (PLS) extracted from all summary statistics.

| Model | Ancestral sources | Layer/s | Marginal density | Wegmann's *p*-value | Bayes factor | *R*<sup>2</sup> |  |  |
| --- | --- | --- | --- | --- | --- | --- | --- | --- |
|  |  |  |  |  |  | *K*<sub>MAX</sub> | *m* | *N*<sub>ANC</sub> |
| *O. bolivari* |  |  |  |  |  |  |  |  |
| A Colonization | LGM<sub>MIROC</sub> | LGM<sub>MIROC</sub> → Current | 9.14 × 10<sup>-07</sup> | 0.368 | 2.09 | 0.78 | 0.61 | 0.92 |
| B Colonization | LGM<sub>CCSM</sub> | LGM<sub>CCSM</sub> → Current | 1.91 × 10<sup>-06</sup> | 0.174 | — | 0.73 | 0.61 | 0.91 |
| C Isolation | Current | Current | 1.52 × 10<sup>-13</sup> | <0.001 | 1.26 × 10<sup>07</sup> | 0.79 | 0.66 | 0.91 |
| D Colonization | LGM<sub>MIROC</sub> | Flat landscape | 2.76 × 10<sup>-09</sup> | 0.006 | 6.93 × 10<sup>02</sup> | 0.69 | 0.68 | 0.93 |
| E Colonization | LGM<sub>CCSM</sub> | Flat landscape | 9.15 × 10<sup>-08</sup> | 0.012 | 2.09 × 10<sup>01</sup> | 0.70 | 0.70 | 0.91 |
| F Isolation | Current | Flat landscape | 3.67 × 10<sup>-13</sup> | <0.001 | 5.22 × 10<sup>06</sup> | 0.69 | 0.72 | 0.90 |
| *O. femoralis* |  |  |  |  |  |  |  |  |
| A Colonization | LGM<sub>MIROC</sub> | LGM<sub>MIROC</sub> → Current | 1.26 × 10<sup>-06</sup> | 0.026 | 1.28 × 10<sup>01</sup> | 0.36 | 0.68 | 0.96 |
| B Colonization | LGM<sub>CCSM</sub> | LGM<sub>CCSM</sub> → Current | 8.82 × 10<sup>-06</sup> | 0.195 | 1.83 | 0.28 | 0.66 | 0.93 |
| C Isolation | Current | Current | 1.00 × 10<sup>-100</sup> | <0.001 | 1.61 × 10<sup>95</sup> | 0.41 | 0.59 | 0.92 |
| D Colonization | LGM<sub>MIROC</sub> | Flat landscape | 6.35 × 10<sup>-09</sup> | 0.001 | 2.54 × 10<sup>03</sup> | 0.24 | 0.69 | 0.96 |
| E Colonization | LGM<sub>CCSM</sub> | Flat landscape | 1.61 × 10<sup>-05</sup> | 0.239 | — | 0.19 | 0.73 | 0.96 |
| F Isolation | Current | Flat landscape | 1.00 × 10<sup>-100</sup> | <0.001 | 1.61 × 10<sup>95</sup> | 0.19 | 0.66 | 0.91 |

$K_{MAX}$ , carrying capacity of the deme with highest suitability (100%)
$m$ , migration rate per deme per generation
$N_{ANC}$ , ancestral population size

i. Colonization of sky islands from glacial refugia (Models A, B, D and E). The first two models (Models A and B) are dynamic models (*sensu* He et al., 2013) incorporating the colonization process from hypothetical glacial refugia and distributional shifts resulted from the interaction between the species bioclimatic envelope and Pleistocene glacial cycles (e.g., He et al., 2013; Massatti & Knowles, 2016). In these scenarios, carrying capacities change over time according to climatic suitability maps obtained from projections of the ENM to the present and the LGM bioclimatic conditions under the MIROC-ESM (Model A) and CCSM (Model B) general atmospheric circulation models (see section *Environmental niche modelling*). These models considered landscapes from three consecutive time periods (LGM, intermediate, current) reflecting temporal shifts in the spatial distribution of environmentally suitable areas for the species in response to climate changes since the LGM (e.g., He et al., 2013; Massatti & Knowles, 2016). Forward demographic simulations under this model initialized 21 ka BP from hypothesized refugial populations (i.e., source populations, each one with an effective population size of *N*_*ANC*_) that were located at every habitat patch predicted as suitable for each focal species during the LGM according to MIROC-ESM (Model A) and CCSM (Model B) projections. Specifically, suitable habitat patches during the LGM were identified as those cells with a probability of presence of the species above the maximum training sensitivity plus specificity (MTSS) logistic threshold for occurrence from MAXENT (Liu et al., 2005). Finally, we generated two more models (Models D and E) analogous to the previous ones but in which carrying capacities (*k*) are homogeneous across space and through time. These static models (*sensu* He et al., 2013) are equivalent to a flat landscape or an isolation-by-distance model and only differ among them in the location of ancestral populations, which were based on the patches of suitable habitat identified under LGM-MIROC (Model D) and LGM-CCSM (Model E) bioclimatic conditions.
ii. Population isolation in sky islands (Models C and F). The first model of this set (Model C) is a static model representing genetic drift associated with the isolation of populations in sky-islands according to the current geographical configuration of suitable habitats. In this model, carrying capacities do not change over time and are defined by an environmental suitability layer obtained from the projection of the ENM to the current bioclimatic conditions. Forward demographic simulations under this model initialized 7 ka BP from hypothesized refugial populations located in every patch of habitat predicted as suitable during present time according to the MTSS logistic threshold for occurrence from MAXENT. Thus, this model hypothesizes that the spatial distribution of contemporary genetic variation reflects population fragmentation and isolation in sky islands since the Mid-Holocene (e.g., Knowles & Massatti, 2017). As done for the first set of models, we also generated a scenario (Model F) analogous to the previous one but in which carrying capacities (*k*) are homogeneous across space (i.e., equivalent to an isolation-by-distance model; He et al., 2013).

#### Model choice and parameter estimation

We used an Approximate Bayesian computation (ABC) framework to perform model selection and parameter estimation, as implemented in ABCTOOLBOX programs (TRANSFORMER and ABCESTIMATOR) and R scripts (*findPLS*) (Wegmann et al., 2010). We used the R package *pls* v.2.6-0 (Mevik & Wehrens, 2007) and the *findPLS* script to extract partial least squares (PLS) components with Box-Cox transformation from the summary statistics of the first 10,000 simulations for each model (Wegmann et al., 2010). The first five PLSs extracted from the summary statistics were used for ABC analyses, as the root-mean-squared error (RMSE) of the three demographic parameters employed (*K*_MAX_, *m, N*_ANC_; see Methods S3 for details) for the two species did not decrease significantly with additional PLSs (Figure S1). We used the linear combinations of summary statistics obtained from the first 10,000 simulations for each model to transform all datasets (observed empirical and simulated datasets) with the program TRANSFORMER (Wegmann et al., 2010). For each model, we retained the 1,000 simulations (0.5%) closest to observed empirical data and used them to approximate marginal densities and posterior distributions of the parameters with a postsampling regression adjustment using the ABC-GLM (general linear model) procedure detailed in Leuenberger & Wegmann (2010) and implemented in ABCESTIMATOR. We used Bayes factors (BF) for model selection (Kass & Raftery, 1995).

#### Model validation

To evaluate the ability of each model to generate the empirical data, we calculated the Wegmann’s *p*-value from the 1,000 retained simulations (Wegmann et al., 2010). We also assessed the potential for a parameter to be correctly estimated by computing the proportion of parameter variance that was explained (i.e., the coefficient of determination, *R*^2^) by the retained PLSs. For the most supported model for each species, we determined the accuracy of parameter estimation using a total of 1,000 pseudo-observation datasets (PODs) generated from prior distributions of the parameters. If the estimation of the parameters is unbiased, posterior quantiles of the parameters obtained from PODs should be uniformly distributed (Wegmann et al., 2010). As with the empirical data, we calculated the posterior quantiles of true parameters for each pseudo run based on the posterior distribution of the regression-adjusted 1,000 simulations closest to each pseudo-observation.

### 2.6. Genetic diversity and historical changes in effective population size

To further explore the processes determining the spatial distribution of genetic variation in the two species we (i) tested the association between genetic diversity and the geographic centrality and habitat patch-size of the studied populations, and (ii) used genomic data to reconstruct changes in effective population size (*N*_*e*_) over time. We employed linear regressions in SPSS v. 26.0 to analyze whether genetic diversity (nucleotide diversity, π) is associated with the distance of each population to the species’ distribution centroid and contemporary suitable habitat patch-size. Given that the precision of genetic diversity estimates may differ among populations due to differences in sample sizes, we used a weighted least square (WLS) method where weight equals the number of genotyped individuals for each population (Table S1). We calculated nucleotide diversity for each population using the program *populations* from STACKS. The centroid of species distribution was calculated in ARCMAP v.10.3 on the basis of polygons (patches) of suitable habitat defined by grid cells with a probability of presence of the focal species above the maximum training sensitivity plus specificity (MTSS) logistic threshold for occurrence from MAXENT (Liu et al., 2005). Similarly, patch-size for each study population was calculated considering the area (in km^2^) of continuous suitable habitat defined as above on the basis of raster maps obtained from projecting ENMs to contemporary bioclimatic conditions.

We inferred changes in effective population sizes (N_e_) over time for each studied population using STAIRWAY PLOT (Liu & Fu, 2015), a flexible multi-epoch model approach based on the site frequency spectrum (SFS) that does not require a predefined demographic model for estimating past demographic histories. To compute a folded SFS for each population, we ran STACKS in order to obtain a VCF file retaining only loci represented in at least 50% of the individuals of the population. To remove all missing data for the calculation of the SFS and minimize errors with allele frequency estimates, each population was down-sampled to five individuals using a custom Python script written by Qixin He and available on Dryad (http://dx.doi.org/10.5061/dryad.23hs1). We ran STAIRWAY PLOT considering a 1-year generation time, assuming the mutation rate per site per generation of 2.8 × 10^−9^ estimated for *Drosophila melanogaster* (Keightley et al., 2014), and performing 200 bootstrap replicates to estimate 95% confidence intervals.

## 3. RESULTS

### 3.1. Genomic data

After quality filtering, 182,588,301 and 74,830,407 reads were retained across all genotyped individuals of *O. bolivari* and *O. femoralis*, respectively (Figure S2). Final exported data sets obtained with STACKS after removing loci that did not meet the different filtering requirements contained 2,262 SNPs for *O. bolivari* and 14,116 SNPs for *O. femoralis*.

### 3.2. Population genetic structure

STRUCTURE analyses showed that log probabilities of the data (LnPr(X|K)) reached a plateau for *K* = 8 in *O. bolivari* and *K*= 5 in *O. femoralis* (Figure S3). The Δ*K* criterion indicated an “optimal” clustering solution for *K* = 5 in *O. bolivari* and *K* = 3 in *O. femoralis* (Figure S3). The two species presented a strong genetic structure, with most (*O. bolivari*) or all (*O. femoralis*) populations being assigned to unique genetic clusters with little or no genetic admixture among them (Figure 2 and S4). In the two species, *K* = 2 separated northwestern (*O. bolivari*: PAND, MAGI, ARAN, and UNIV; *O. femoralis*: ALME and CAZO) from southeastern populations (*O. bolivari*: POZO, BAZA, FILA, RAGU, LUMA, LUJA, and GADO; *O. femoralis*: POYO, MARI, and ESPU). These two main clusters split hierarchically at increasingly higher *K* values, generally grouping the different populations in accordance with their geographical location. Remarkably, populations located in the same “sky island” did not always cluster together at different hierarchical levels. For instance, in *O. bolivari*, population UNIV grouped with northwestern populations (PAND, MAGI, and ARAN) rather than with other populations (LOMA and RAGU) located within its same mountain range (i.e., the main Sierra Nevada massif; Figure 1a and 2b). In the same line, population POYO from *O. femoralis* clustered together with the highly isolated southeastern populations (MARI and ESPU) rather than with the two other populations (ALME and CAZO) located within the Sierra de Cazorla massif and with which it is expected to be currently more connected (Figure 1b and 2b). Principal component analyses for both species yielded analogous results to those obtained with STRUCTURE, separating along PC1 northwestern from southeastern populations and grouping populations according to their geographical location (Figure S5).

**FIGURE 2.**
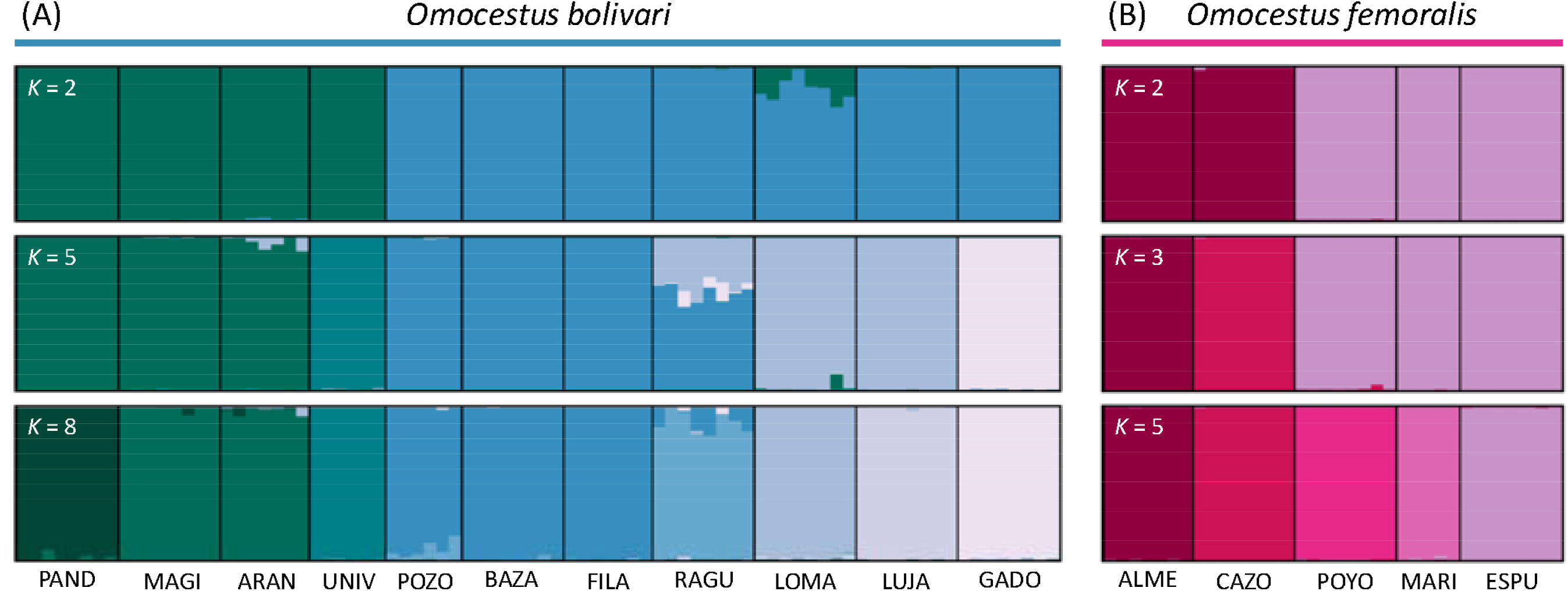
Genetic assignment of individuals based on the results of STRUCTURE for (A) *Omocestus bolivari* (*K* = 2, *K* = 5 and *K* = 8) and (B) *O. femoralis* (*K* = 2, *K* = 3 and *K* = 5). Individuals are partitioned into K colored segments representing the probability of belonging to the cluster with that color. Thin vertical black lines separate individuals from different populations. Population codes as in Table S1.

### 3.3. Environmental niche modelling

We generated final ENMs using the feature class (FC) combination (*O. bolivari* = LQ; *O. femoralis* = LQ) and regularization multiplier (*O. bolivari* = 6.0; *O. femoralis* = 2.0) that minimized AICc across the set of models tested with ENMEVAL for each species. After removing highly correlated variables (*r* > 0.9) and those with a zero percent contribution, our models retained four bioclimatic variables for *O. bolivari* (sorted by percent contribution, BIO8: 36.8 %; BIO7: 34.8 %; BIO11: 16.5 %; BIO2: 11.9 %) and eight variables for *O. femoralis* (BIO11: 37.9 %; BIO2: 33.9 %; BIO7: 13.6 %; BIO8: 7.4 %; BIO19: 4.3 %; BIO15: 1.3 %; BIO12: 1.2 %; BIO18: 0.4 %). Inspection of predicted distributions for the present confirmed that ENMs yielded distribution patterns coherent with the observed current distribution of the two species (Figure 1c-d). However, current distribution maps for the two species also identified as suitable some high elevation areas outside their respective distribution ranges. In the two species, over-predicted areas included partially (*O. bolivari*) or totally (*O. femoralis*) the distribution range of each other, which is not surprising given their similar habitat, elevation, and environmental requirements. Projections of present-day climate niche envelopes to LGM climatic conditions under two general atmospheric circulation models (CCSM4 and MIROC-ESM) suggest that the two species have experienced important distributional shifts in response to Pleistocene glaciations (Figure 1e-h). As expected for montane-alpine organisms, the distribution of the two species expanded into lowland areas during the LGM (Figure 1e-h). However, projections under the two circulation models also presented important differences. Populations of the two species were predicted to be much more connected during the LGM under the MIROC-ESM than under the CCSM model. According to MIROC-ESM, most currently isolated populations of the two species (with the exception of Sierra de Lújar and Sierra de Gádor in *O. bolivari*) became connected by corridors of suitable habitat during the LGM. However, in *O. bolivari*, most contemporary sky islands were also predicted to be isolated from each other during the LGM according to the CCSM model (with the exception of Sierra de Arana and Sierra Nevada; Figure 1g). In the case of *O. femoralis*, the south-westernmost portion of its current distribution was predicted as unsuitable under the CCSM model, suggesting that this species might have experienced important distributional changes beyond elevational shifts (Figure 1h).

### 3.4. Testing alternative demographic models

Based on marginal densities calculated from the 0.5% of retained simulations, the best ranked models in the two species were those incorporating the colonization process from hypothetical glacial refugia (Models A, B, D and E; Table 1). In contrast, models of isolation in contemporary sky islands had considerably lower marginal densities and a difference in Bayes factors > 10^6^ with the most supported model (Table 1), indicating strong relative support for colonization models in the two species (Kass & Raftery, 1995). Moreover, only colonization models were able to simulate genetic data comparable with empirical data (Models A and B in *O. bolivari* and Models B and E in *O. femoralis*), unlike the isolation models in which there was a substantial difference between the likelihoods of the simulated data compared with the empirical data (Wegmann’s *p*-values < 0.001 in all cases; Table 1). Specifically, for *O. bolivari*, dynamic colonization models incorporating changes in habitat suitability over time under both the CCSM and MIROC projections (Models A and B) were those most supported by our empirical data and small Bayes factors (BF < 3; Table 1) suggest that they are statistically indistinguishable (Kass & Raftery, 1995). In *O. femoralis*, both dynamic and static (i.e., flat landscape) colonization models based on CCSM projections (Models B and E) were the most supported by our empirical data and statistically indistinguishable (BF < 2; Table 1). The dynamic colonization model based on MIROC bioclimatic conditions also had a low Bayes factor (< 20), but its low Wegmann’s *p*-value (< 0.03) indicates that it was not capable of generating data compatible with our empirical data (Table 1).

Posterior distributions of parameters under the most probable models were considerably distinct from the prior, indicating that the simulated data contained information relevant to estimating the parameters (Figure S6). Comparison of the posterior distributions before and after the ABC-GLM also showed the improvement that this procedure had on parameter estimates (Figure S6). The posterior distribution of *K*_MAX_ in *O. femoralis* was flat, indicating uncertainty in the estimation of this parameter (Figure 6Sb). Accordingly, the coefficients of determination (*R*^2^) from a multiple regression between each demographic parameter and the five retained PLS indicated that the employed summary statistics had a moderately high potential to correctly estimate all the parameters except *K*_MAX_ in *O. femoralis* (Table 1). Posterior quantiles of *m* in *O. bolivari, K*_MAX_ in *O. femoralis* and *N*_ANC_ in both taxa significantly deviated from a uniform distribution, indicating a potential bias in the estimation of these parameters (Figure S7).

### 3.5. Genetic diversity and historical changes in effective population sizes

Visual inspection of the spatial distribution of genetic variation in the two species evidenced that peripheral populations and those located in smaller “sky islands” tended to present lower levels of genetic diversity (Figure 1a, b). Accordingly, genetic diversity (π) in populations of *O. bolivari* was negatively associated with population centrality (*r* = −0.913, *t* = −6.72, *P* < 0.001; Figure 3a) and showed a positive relationship with patch size (*r* = 0.733, *t* = 3.23, *P* = 0.010; Figure 3b). In the case of *O. femoralis*, genetic diversity was positively associated with patch size (*r* = 0.878, *t* = 3.17, *P* = 0.050; Figure 3d) but the negative relationship with population centrality did not reach statistical significance (*r* = 0.743, *t* = −1.92, *P* = 0.150; Figure 3c). STAIRWAY PLOT analyses for *O. bolivari* revealed similar results across most of the populations, with an abrupt demographic expansion led by the advance of the glaciation till its maximum level around 21 ka BP (Figure 4a, b). Afterwards, populations experienced a decline in *N*_*e*_ until recent times (Figure 4a, b). One exception was the highly isolated populations PAND, MAGI and LUJA, which presented a general demographic stability during the last 100 ka (Figure 4b). In the case of *O. femoralis*, most populations showed idiosyncratic demographic histories, which ranged from expansions during the last glacial period followed by reductions in *N*_e_ at the onset of the Holocene (ALME and MARI) to marked bottlenecks around the LGM (ESPU and CAZO) (Figure 4c, d).

**FIGURE 3.**
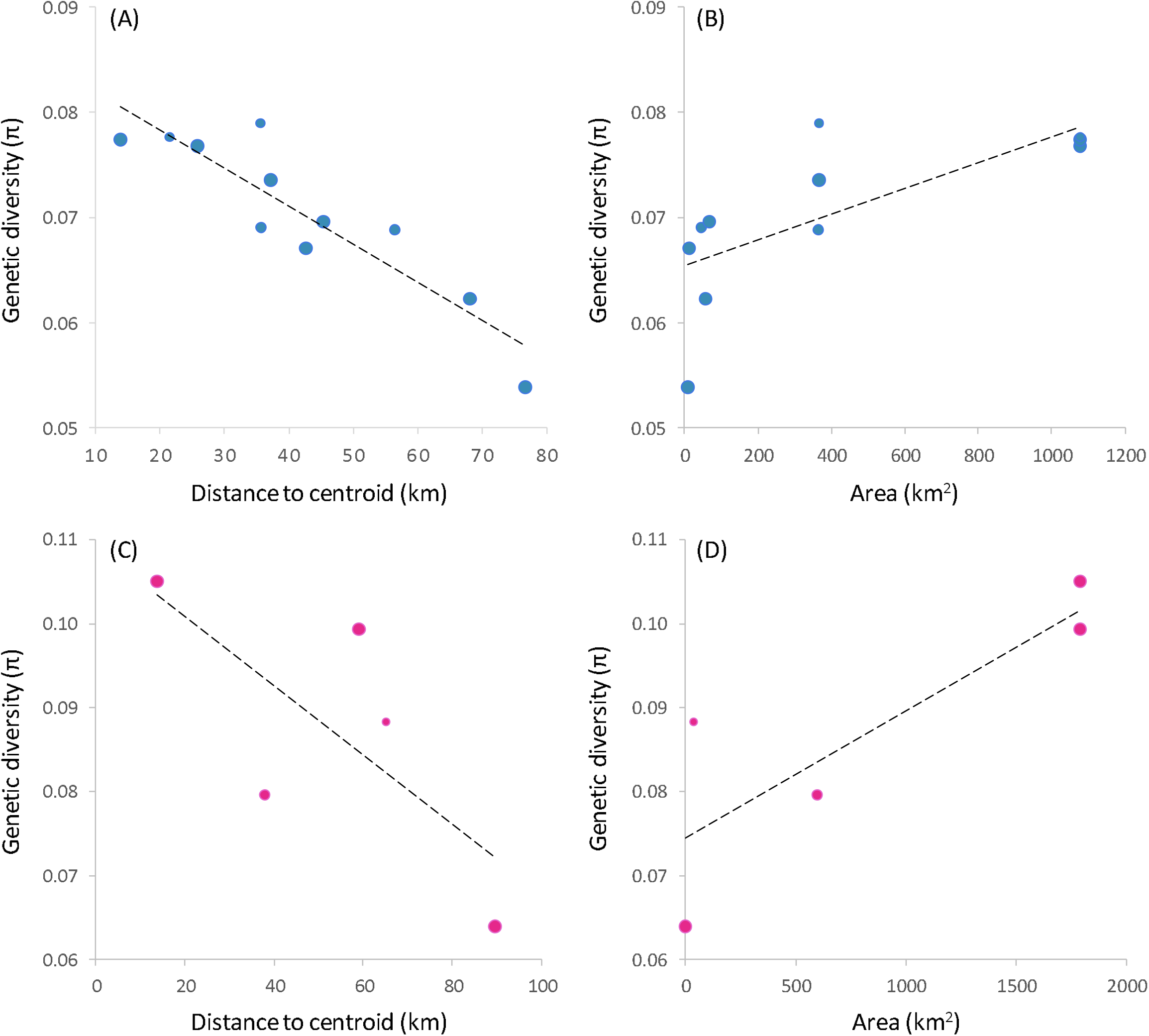
Relationship between the genetic diversity (π) of populations and (A, C) distance to centroid of species’ distribution and (B, D) area of climatically suitable habitat for (A, B) *Omocestus bolivari* and (C, D) *O. femoralis*. Regression lines are indicated and dot size is proportional to sample size for each studied population.

**FIGURE 4.**
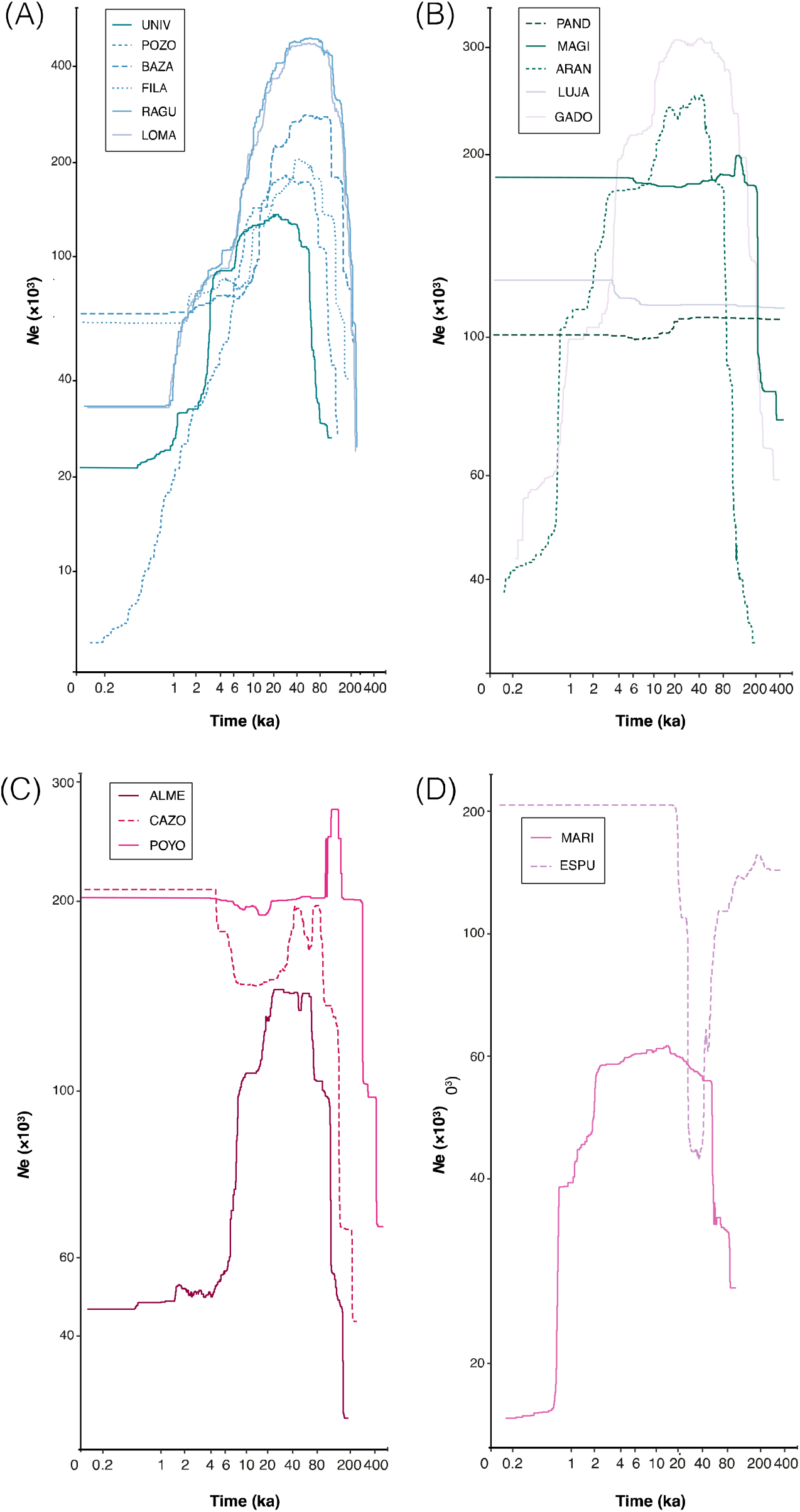
Demographic history of each studied population of (A, B) *O. bolivari* and (C, D) O. *femoralis* inferred using STAIRWAY PLOT. Lines show the median estimate of effective population size (*N*_*e*_) over time (ka) for populations located in (A, C) large and (B, D) small sky islands. Population codes shown in the legend of each panel are described in Table S1.

## 4. DISCUSSION

Our study contributed to a better understanding of the demographic processes underlying spatial patterns of genetic variation in cold-adapted biotas currently forming severely fragmented populations in sky-island archipelagos from temperate regions (Hewitt, 1996; Knowles & Massatti, 2017; Perrigo et al., 2020). Specifically, the combination of population genomic data, niche modelling and spatiotemporally explicit coalescent-based simulations supported population connectivity during glacial periods followed by colonization and population isolation in sky islands as the most supported scenario explaining contemporary patterns of population genetic diversity and structure in the two focal taxa. Although our analyses indicate that post-glacial fragmentation and genetic drift have not blurred the genomic signatures left by historical patterns of population connectivity, the lower levels of genetic diversity in peripheral populations confined to small patches of suitable habitat (i.e., small-sized sky islands) evidence the genetic consequences of long-term isolation. This study exemplifies how the integration of multiple lines of evidence provided by a comprehensive suite of analytical approaches applied to two independent spatial and biological replicates can help to identify concerted or idiosyncratic evolutionary and demographic histories of species with similar habitat requirements and, ultimately, reach more general conclusions about the way organisms and whole communities have responded to Quaternary climatic oscillations. These aspects are of pivotal importance to understand the dynamics of montane and alpine ecosystems from mid and low latitude regions, which not only harbor high rates of local endemism but have been also identified among the most vulnerable to extinction due to ongoing climate warming and the progressive shrink of their habitats (Perrigo et al., 2020).

### 4.1. Marked genetic structure in sky islands

The contemporary genetic structure of *O. bolivari* and *O. femoralis* exhibited many similarities, with both taxa showing a marked northwest-southeast pattern of genetic differentiation and most genetic clusters identified at the different hierarchical levels presenting a very low degree of genetic admixture (Figure 2). The northwest-southeast pattern of differentiation likely reflects the predominant connectivity and origin of source populations that led to the colonization of sky islands from the more widespread lowland distributions presented by the two species during glacial periods, as supported by projections of environmental niche models to LGM bioclimatic conditions under alternative general circulation models (Figure 1). Remarkably, the two taxa presented populations that, being located on the same mountain range, are assigned to different genetic clusters, which, in turn, are shared with populations located in different sky-islands. This is exemplified by the genetic assignment of UNIV and POYO populations from *O. bolivari* and *O. femoralis*, respectively. Population UNIV is situated in a slope facing north in Sierra Nevada massif but shares a common ancestry with the northernmost populations of the species rather than with the other two populations (LOMA and RAGU) located within the same mountain range (Figure 2a). Similarly, the large Sierra de Cazorla massif encompasses populations CAZO, ALME and POYO, but the latter clusters with the highly isolated and southernmost populations MARI and ESPU (Figure 2b). This indicates that patterns of genetic structure at different hierarchical levels are in some instances more congruent with the geographical arrangement of populations than with their location in a specific sky island. Projections of ENMs to LGM bioclimatic conditions point to the connection of these populations during cold periods, indicating that apparently incongruent patterns of genetic structure likely reflect the identity of ancestral glacial-source populations rather than the spatial configuration of contemporary suitable habitats and hypothetical corridors to gene flow. The low dispersal capacity of the brachypterous focal taxa (Presa et al., 2016a, b), the considerable topographic complexity (i.e., ridges and steep slopes) within the main massifs (> 3,400 m in Sierra Nevada) of the region, and the limited ability of grasshoppers to move across abrupt landscapes (e.g., González-Serna et al., 2019; Tonzo et al., 2019) might collectively explain the deep genetic structure and low or absent genetic admixture among populations located within the same sky island even if these are predicted to be nowadays connected by corridors of suitable habitat according to ENMs.

### 4.2. Post-glacial colonization of sky-islands

Model selection under an ABC framework revealed that the spatially-explicit demographic scenarios most supported by empirical data were those in which the spatial distribution of contemporary genomic variation was shaped by processes of post-glacial colonization of sky islands from source populations defined by the spatial location of environmentally suitable areas during the last ice age (Knowles & Massatti, 2017; Mouret et al., 2011). This is in agreement with the classical pattern of glacial expansion and interglacial contraction suggested for montane species from temperate regions (e.g., DeChaine & Martin, 2005). In contrast, models of isolation in sky islands since the onset of the Holocene had a very low statistical support, indicating that genetic drift after post-glacial fragmentation cannot solely explain the genetic makeup of contemporary populations. Although scenarios based on colonization of sky-islands were the only ones able to simulate genomic data compatible with observed data, our approach was not able to distinguish between dynamic models considering changes in landscape heterogeneity since the LGM and static models considering a diffusion scenario of expansion from glacial source populations (see also Knowles & Massatti, 2017). These results are surprising considering that the two focal taxa have strict habitat preferences for high elevation areas and unsuitable lowlands are expected to act as impassable barriers to dispersal, as evidenced by distribution models (Figure 1) and strong genetic structure of contemporary populations (Figure 2). Simple Mantel tests indicate that genetic differentiation (*F*_ST_) is positively correlated with geographical distances among populations in *O. bolivari* (*r* = 0.57, *P* < 0.001, *n* = 11) and marginally in *O. femoralis* (*r* = 0.60, *P* = 0.058, *n* = 5). Processes of genetic admixture and drift mediated by population expansion and contraction during Quaternary glacial cycles is expected to have resulted in this pattern of isolation-by-distance (Slatkin, 1993), which is similar to the expected outcome of migration-genetic drift equilibrium under a static, flat landscape scenario (He et al., 2013; Knowles & Massatti, 2017). Thus, admixture among nearby populations is likely to have reduced the power of our analyses to discriminate between dynamic (Models A-B) and static (Models D-E) scenarios of postglacial colonization, particularly in *O. femoralis* for which only five populations were available for analyses (Table S1).

The two general circulation palaeoclimatic models supported the expansion of the two focal taxa toward lowland areas during the LGM, but their respective populations were predicted to be much more connected under the MIROC-ESM than under the CCSM model (Figure 1). Although differences between palaeoclimatic models in predicting past species distributions is a frequent outcome of most studies (e.g., Ramírez-Barahona & Eguiarte, 2014), the relative support for contrasting predictions is very rarely validated using independent sources of information (Nogués-Bravo, 2009). Such differences seemed to be captured by our model-based approach, which revealed that glacial-source populations identified for the two focal taxa on the basis of projections of ENMs to bioclimatic conditions under the CCSM general circulation model tended to provide a better fit to our genomic data than those obtained based on MIROC projections (BF > 2 in all cases; Table 1). Although differences in the relative likelihood for scenarios based on the two palaeoclimatic models are small and must be interpreted with extreme caution, our results highlight the potential of spatiotemporally explicit coalescent-based simulations to refine hypotheses about the location of glacial refugial (Bemmels et al., 2019) and validate distributional shifts inferred from ENMs that cannot usually be contrasted with independent sources of information (e.g., fossil records; Fordham et al., 2014; Metcalf et al., 2014).

### 4.3. Genetic diversity and demographic reconstructions

The spatial distribution of genetic variation in the two species evidenced that peripheral populations and those located in smaller sky islands tend to present comparatively lower levels of genetic diversity (Figure 1). These findings are consistent with the center-periphery model, which predicts that populations located on the species range edges become progressively smaller, are spatially more isolated, and harbor lower levels of genetic diversity than populations situated at the core of the distribution (e.g., Lira-Noriega & Manthey, 2014). In contrast, genomic-based demographic reconstructions indicate that, in general terms, most populations presented larger effective population sizes during the last glacial period and experienced demographic declines at the onset of the Holocene (Figure 4), in agreement with palaeoclimatic-based reconstructions of species distributions (Figure 1). Inference of changes of *N*_*e*_ through time did not reveal obvious differences between populations located in large vs. small sky islands, suggesting that contemporary populations, and their ancestral sources, reacted in a similar fashion to Quaternary climatic oscillations independently of their current geographical location. This is not surprising given the small range of the two focal taxa (Presa et al., 2016a, b), with most distant populations separated by <150 km, and the likely similar regional impact of Pleistocene glacial cycles across the entire distribution of the species. Also, it must be considered that, according to our own field observations, small habitat patches can often sustain high local densities of the species, which might contribute to maintain effective population sizes above a certain threshold and avoid strong genetic drift even in populations currently confined to tiny sky islands. Collectively, these results indicate that although the genetic makeup of contemporary populations has been shaped in a great extent by historical processes of genetic admixture (glacial periods) and drift during the retreat to inter-glacial refugia (e.g., Knowles & Massatti, 2017), contemporary isolation has also contributed to erode the levels of genetic diversity of populations persisting in peripheral sky islands of small size (Lira-Noriega & Manthey, 2014; Rubidge et al., 2012).

## 5. CONCLUSIONS

By considering two biological replicates from the sky island archipelago of the Baetic System, our study highlights the potential of integrating different sources of information to infer the evolutionary and demographic processes shaping spatial patterns of genetic variation in cold-adapted faunas from temperate regions. Our spatiotemporally explicit simulations and testing of alternative demographic scenarios supported population expansion during cold periods and subsequent postglacial colonization and isolation in sky islands as the most likely explanation for the current distribution of genetic variation in both *O. bolivari* and *O. femoralis*. Global warming is expected to threat the persistence of alpine and cold adapted organisms from mid and low latitude regions by reducing the availability of suitable habitats confined to mountain tops (Rubidge et al., 2012). The similar demographic responses inferred for the two focal taxa to past climate changes suggest that they and other co-distributed organisms with similar life-history traits will probably present concerted responses to ongoing global change. Thus, our results can help to establish unified conservation strategies aimed at preserving the extraordinary rates of local endemism of this and other mountain biodiversity hotspots, from a community-level perspective (Perrigo et al., 2020).

## Acknowledgements

We are much indebted to Anna Papadopoulou for her valuable help in study design and useful comments, suggestions and corrections on a first draft of the manuscript. We also wish to thank to Pedro J. Cordero and Víctor Noguerales for their help during fieldwork and providing samples from some populations, Amparo Hidalgo for support during laboratory work, and Sergio Pereira (The Centre for Applied Genomics) for Illumina sequencing. Logistical support was provided by Laboratorio de Ecología Molecular (LEM-EBD) and Laboratorio de Sistemas de Información Geográfica y Teledetección (LAST–EBD) from Estación Biológica de Doñana. We also thank to Centro de Supercomputación de Galicia (CESGA) and Doñana’s Singular Scientific-Technical Infrastructure (ICTS-RBD) for access to computer resources. This study was funded by the Spanish Ministry of Economy and Competitiveness and the European Regional Development Fund (ERDF) (CGL2014-54671-P and CGL2017-83433-P). VT was supported by an FPI predoctoral fellowship (BES-2015-73159) from Ministerio de Economía y Competitividad.

## DATA AVAILABILITY STATEMENT

Raw Illumina reads have been deposited at the NCBI Sequence Read Archive (SRA) under BioProject XXXXX. Input files for all analyses are available for download on Figshare (https://doi.org/XXXXX).

## SUPPORTING INFORMATION

Additional supporting information may be found online in the Supporting Information section.

## Notes

### Competing Interest Statement

The authors have declared no competing interest.

