## Supporting information for "Glacial connectivity and current population fragmentation in sky-islands explain the contemporary distribution of genomic variation in two narrow-endemic montane grasshoppers from a biodiversity hotspot"

**Contents:**

**Supplemental methods**

**Methods S1** Processing of genomic data

**Methods S2** Environmental niche modelling

**Methods S3** Testing alternative demographic models

**Supplemental TABLES**

**TABLE S1** Geographical location and genetic diversity of studied populations

**Supplemental figures**

**Figure S1** Root Mean Square Error (RMSE) of parameter estimates

**Figure S2** Number of reads before and after different quality filtering steps by stacks

**Figure S3** Log probability of the data and the magnitude of Δ*K* for Bayesian clustering analyses in structure

**Figure S4** Genetic assignment of individuals based on the results of Bayesian clustering analyses in structure

**Figure S5** Principal component analysis (PCAs) of genetic variation

**FIGURE S6** Posterior distribution of parameter estimates for the most supported models

**Figure S7** Distribution of posterior quantiles of true parameter values based on pseudo-observed data sets

**References**

**Supplemental methods**

**Methods S1** Processing of genomic data

We used the different programs distributed as part of the stacks v. 1.35 pipeline (*process_radtags*, *ustacks*, *cstacks*, *sstacks*, and *populations*) to assemble our sequences into *de novo* loci and call genotypes (Catchen et al., 2011, 2013; Hohenlohe et al., 2010). We demultiplexed and filtered reads for overall quality using the program *process_radtags*, retaining reads with a Phred score > 10 (using a sliding window of 15%), no adaptor contamination, and that had an unambiguous barcode and restriction cut site. We screened raw reads for quality with fastqc v. 0.11.5 (<http://www.bioinformatics.babraham.ac.uk/projects/fastqc/>) and trimmed all sequences to 129-bp using seqtk (Heng Li, <https://github.com/lh3/seqtk>) in order to remove low-quality reads near the 3´ ends. We assembled filtered reads *de novo* into putative loci with the program *ustacks*. We set the minimum stack depth (*m*) to three and allowed a maximum distance of two nucleotide mismatches (*M*) to group reads into a “stack”. We used the “removal” (*r*) and “deleveraging” (*d*) algorithms to eliminate highly repetitive stacks and resolve over-merged loci, respectively. We identified single nucleotide polymorphisms (SNPs) at each locus and called genotypes using a multinomial-based likelihood model that accounts for sequencing errors, with the upper bound of the error rate (*ε*) set to 0.2 (Catchen et al., 2011, 2013; Hohenlohe et al., 2010). Then, we built a catalogue of loci using the *cstacks* program, with loci recognized as homologous across individuals if the number of nucleotide mismatches between consensus sequences (*n*) was ≤2. Finally, we matched each individual data against this catalogue using the program *sstacks* and exported output files in different formats for subsequent analyses using the program *populations*. We used the “blacklist” option to remove outlier loci (i.e., putatively under selection) identified using bayescan v.2.1 (Foll & Gaggiotti, 2008) and the hierarchical and non-hierarchical island models (FDIST method; Excoffier et al., 2009) implemented in arlequin v. 3.5.2.2 (Excoffier & Lischer, 2010) (see Ortego et al., 2018 for details). For all downstream analyses, we exported only the first SNP per RAD locus and retained loci with a minimum stack depth ≥ 5 (*m* = 5), a minimum minor allele frequency (MAF) ≥ 0.01 (*min_maf* = 0.01) and that were represented in all populations (*p* = 11 for *O. bolivari* and *p* = 5 for *O. femoralis*) and the 50% of the individuals within each population (*r* = 0.5).

**Methods S2** **Environmental niche modelling**

To build ENMs, we used the maximum entropy algorithm implemented in maxent v.3.3.3 (Phillips et al., 2006; Phillips & Dudik, 2008) and the 19 bioclimatic variables from the worldclim dataset (http://www.worldclim.org/) interpolated to 30-arcsec resolution (~1 km^2^ cell size) (Hijmans et al., 2005). To generate climate suitability maps during the LGM, we projected the ENM onto LGM bioclimatic conditions derived from the CCSM4 (Community Climate System Model; Braconnot et al., 2007) and the MIROC-ESM (Model of Interdisciplinary Research on Climate; Hasumi & Emori, 2004) general atmospheric circulation models. We built ENMs using our own species occurrence data and records available in the literature (Prunier, 2014 and references therein). Prior to modelling, we mapped and examined all records to identify and exclude those having obvious geo-referencing errors. After data filtering and excluding duplicates (i.e., records falling within the same grid cell), we retained a total of 26 occurrence records for *O. bolivari* and 16 records for *O. femoralis*. We used the r package enmeval (Muscarella et al., 2014) and the Akaike’s Information Criterion corrected for small sample size (AICc) (Warren & Seifert, 2011) to conduct parameter tuning and determine the optimal feature class (FC) and regularization multiplier (RM) settings for maxent. We tested a total of 248 models of varying complexity by combining a range of regularization multipliers (RM) (from 0 to 15 in increments of 0.5) with eight different feature classes (FC) combinations (L, LQ, LQP, H, T, LQH, LQHP, LQHPT, where L = linear, Q = quadratic, H = hinge, P = product and T = threshold) and followed the approach detailed in González-Serna et al. (2019) for variable selection.

**Methods S3** Testing alternative demographic models

We used the integrative distributional, demographic and coalescent (iDDC) approach (He, Edwards, & Knowles, 2013) and an Approximate Bayesian Computation (ABC) framework (Beaumont et al., 2002; Csilléry et al., 2010) to test alternative scenarios representing different hypotheses about how landscape heterogeneity (or its lack thereof) and colonization from glacial refugia *vs*. contemporary population isolation in sky islands explain the spatial distribution of genomic variation in our two focal species. This approach is described in He et al. (2013) and consists of three main steps: (i) constructing alternative demographic models representing different hypotheses about the processes shaping spatial patterns of genetic structure and diversity; (ii) running demographic and genetic simulations under each model using the software splatche2 (see Ray et al., 2010); (iii) evaluating the fit of observed genomic data (i.e., empirical genomic data obtained after genotyping sampled populations) to the genetic expectations under each model, identifying the most probable model/s, and estimating demographic parameters (e.g., González-Serna et al., 2019; He et al., 2013). Below we describe the most relevant aspects of this approach.

*Constructing demographic models*. We generated two main sets of models that differ in the hypothetical demographic processes that have shaped spatial patterns of contemporary genetic variation (Table 1):

i) Colonization of sky islands from glacial refugia (Models A, B, D and E). The first two models (Models A and B) are dynamic models (*sensu* He et al., 2013) incorporating the colonization process from hypothetical glacial refugia and distributional shifts resulted from the interaction between the species bioclimatic envelope and Pleistocene glacial cycles (e.g., He et al., 2013; Massatti & Knowles, 2016). In these scenarios, carrying capacities change over time according to climatic suitability maps obtained from projections of the ENM to the present and the LGM bioclimatic conditions under the MIROC-ESM (Model A) and CCSM (Model B) general atmospheric circulation models (see section *Environmental niche modelling*). These models considered landscapes from three consecutive time periods (LGM, intermediate, current) reflecting temporal shifts in the spatial distribution of environmentally suitable areas for the species in response to climate changes since the LGM (e.g., He et al., 2013; Massatti & Knowles, 2016). As done in previous studies, carrying capacities were scaled proportionally to logistic climatic suitability scores obtained from the ENM for each time period (e.g., He et al., 2013; Knowles & Massatti, 2017; Massatti & Knowles, 2016). Thus, we assumed that the carrying capacity for each grid cell was proportional to the estimated probability of presence of the species in that grid cell. As splatche2 requires a single raster file with positive integer numbers, we used arcmap v.10.3 to categorize cell values from logistic ENM maps (ranging continuously from 0 to 1) under each time period into 20 bins of equal magnitude (i.e., intervals of 0.05) (e.g., Bemmels et al., 2016). Then, we ran a custom Python script written by Q. He (deposited in Dryad; Bemmels et al., 2016) to convert the maps from the different time periods into a single raster map in which each category represents a unique combination of climatic suitability bins across the three time periods (e.g., Bemmels et al., 2016; He et al., 2013; Massatti & Knowles, 2016). Climatic suitability bins corresponding to each of the three periods (LGM, intermediate, current) were applied to one-third of the total number of simulated generations. Forward demographic simulations under this model initialized 21 ka BP from hypothesized refugial populations (i.e., source populations, each one with an effective population size of *N*_ANC_) that were located at every habitat patch predicted as suitable for each focal species during the LGM according to MIROC-ESM (Model A) and CCSM (Model B) projections. Specifically, suitable habitat patches during the LGM were identified as those cells with a probability of presence of the species above the maximum training sensitivity plus specificity (MTSS) logistic threshold for occurrence from maxent (Liu et al., 2005). We allowed source population overflow, so that all individuals exceeding the carrying capacity of initial populations (i.e., demes) spread around neighboring cells (Ray et al., 2010). Finally, we generated two more models (Models D and E) analogous to the previous ones but in which carrying capacities (*k*) are homogeneous across space and through time. These static models (*sensu* He et al., 2013) are equivalent to a flat landscape or an isolation-by-distance model and only differ among them in the location of ancestral populations, which were based on the patches of suitable habitat identified under LGM-MIROC (Model D) and LGM-CCSM (Model E) bioclimatic conditions.

ii) Population isolation in sky islands (Models C and F). The first model of this set (Model C) is a static model representing genetic drift associated with the isolation of populations in sky-islands according to the current geographical configuration of suitable habitats. In this model, carrying capacities do not change over time and are defined by an environmental suitability layer obtained from the projection of the ENM to the current bioclimatic conditions. Forward demographic simulations under this model initialized 7 ka BP from hypothesized refugial populations located in every patch of habitat predicted as suitable during present time according to the MTSS logistic threshold for occurrence from maxent. Thus, this model hypothesizes that the spatial distribution of contemporary genetic variation reflects population fragmentation and isolation in sky islands since the Mid-Holocene (e.g., Knowles & Massatti, 2017). As done for the first set of models, we also generated a scenario (Model F) analogous to the previous one but in which carrying capacities (*k*) are homogeneous across space (i.e., equivalent to an isolation-by-distance model; He et al., 2013).

*Demographic and genetic simulations*. We used splatche2 to perform forward-in-time demographic simulations followed by backward-in-time genetic (coalescent) simulations under each model (see Ray et al., 2010), which are expected to produce contrasting patterns of genetic variation due to differences among scenarios in the location of ancestral populations and the way carrying capacities vary across the landscape and through time (see Massatti & Knowles, 2016). To have a computationally tractable number of cells for demographic simulations, we statistically downscaled cell sizes to 2-arcminute (~4 km^2^) (e.g., Bemmels et al., 2016; Massatti & Knowles, 2016). Despite of the studied species having a generation time of one year, we scaled it by a factor of 15 to make simulations computationally tractable (e.g., Massatti & Knowles, 2016), which results in a total of 1,400 generations from the LGM to present (21 ka) and 467 generations from the mid-Holocene to present (7 ka). Because of this scaling, any biological interpretation of absolute values of population genetic parameters would need to be adjusted accordingly (Massatti & Knowles, 2016). For each model, we ran 200,000 simulations using the uniform priors for the three demographic parameters of the spatially explicit coalescent: carrying capacity of the deme with highest suitability (*K*_max_; range of log(*K*_max_) for *O. bolivari*: 4.3, 6.3; range of log(*K*_max_) for *O. femoralis*: 5.2, 7.2), migration rate per deme per generation (*m*; range of log(*m*) for *O. bolivari*: -3.1, 2.0; range of log(*m*) for *O. femoralis*: -2.3, -1.2), and ancestral population size (*N*_ANC_; range of log(*N*_ANC_) for *O. bolivari*: 1.7, 3.7; range of log(*N*_ANC_) for *O. femoralis*: 3.5, 5.5). Before setting the final prior values used in the simulations for each species, we tested a broad range of priors in pilot runs in order to identify those that result in the colonization of the landscape and generate genetic data within the range of observed empirical data. Following each time-forward demographic simulation, a spatially-explicit time-backward coalescent model informed by the deme-specific demographic parameters (*K*, *m* and *N*_ANC_) was used to generate genetic data (Currat et al., 2004; Ray et al., 2010). We ran an independent coalescent process to trace the genealogy for each locus from the present to the onset of population expansion from ancestral source populations and beyond, until alleles coalesced in a single ancestral population of size *N*_ANC_. We set a maximum of 10^6^ generations for providing ample time for coalescence. To make simulations computationally tractable, we randomly selected 1,250 loci. Simulated datasets were sampled from the same geographical locations (grid cells) from which the empirical genomic data were obtained (Table S1) and consisted of the same number of loci, number of individuals, and amount and pattern of missing data as in the observed empirical dataset (see Massatti & Knowles, 2016). We used arlsumstat v.3.5.2 (Excoffier & Lischer, 2010) to calculate summary statistics (SS) for simulated datasets under each model, including mean heterozygosity across loci for each population and across populations (*H*), number of segregating sites for each population and across populations (*S*), and pairwise population *F*_ST_ values (79 SS for the eleven populations of *O. bolivari* and 22 SS for the five populations of *O. femoralis*). The same SS were extracted from observed empirical data for each species. We ran all simulations on the high-performance computing cluster from Centro de Supercomputación de Galicia (CESGA, Spain). Simulations required ~7,000 CPU hours per model.

*Model choice and parameter estimation*. We used an Approximate Bayesian computation (ABC) framework to perform model selection and parameter estimation (for an overview of ABC, see Beaumont et al., 2002), as implemented in abctoolbox programs (transformer and abcestimator) and r scripts (*findPLS*) (Wegmann et al., 2010). In order to account for correlations between summary statistics and reduce the “curse of dimensionality” associated with using a large number of statistics (Boulesteix & Strimmer, 2007), we used the r package *pls* v.2.6-0 (Mevik & Wehrens, 2007) and the *findPLS* script to extract partial least squares (PLS) components with Box-Cox transformation from the summary statistics of the first 10,000 simulations for each model (Boulesteix & Strimmer, 2007; Wegmann et al., 2010). The first five PLSs extracted from the summary statistics were used for ABC analyses, as the root-mean-squared error (RMSE) of the three demographic parameters (*K*_MAX_, *m*, *N*_ANC_) for the two species did not decrease significantly with additional PLSs (Figure S1). We used the linear combinations of summary statistics obtained from the first 10,000 simulations for each model to transform all datasets (observed empirical and simulated datasets) with the program transformer (for details about this procedure, see Wegmann et al., 2010). For each model, we retained the 1,000 simulations (0.5%) closest to observed empirical data and used them to approximate marginal densities and posterior distributions of the parameters with a postsampling regression adjustment using the ABC-GLM (general linear model) procedure detailed in Leuenberger & Wegmann (2010) and implemented in abcestimator (see also Csilléry et al., 2010). We used Bayes factors (BF) for model selection, defined as the ratio between marginal densities of the model with the highest marginal density and the alternative model (Jeffreys, 1961). The higher the ratio is, the more supported the first model is. A BF > 20 indicates strong relative support for the first model, while those >150 indicate very strong support (Jeffreys, 1961; Kass & Raftery, 1995; Leuenberger & Wegmann, 2010).

*Model validation*. To evaluate the ability of each model to generate the empirical data, we calculated the Wegmann’s *p*-value from the 1,000 retained simulations (Wegmann et al., 2010). The Wegmann’s *p*-value is calculated as the fraction of the retained simulations with a smaller or equal likelihood than the empirical data, with low values indicating that a model is highly unlikely (Wegmann et al., 2010). We also assessed the potential for a parameter to be correctly estimated by computing the proportion of parameter variance that was explained (i.e., the coefficient of determination, *R*^2^) by the retained PLSs (Neuenschwander et al., 2008). For the most supported model for each species, we determined the accuracy of parameter estimation using a total of 1,000 pseudo-observation datasets (PODs) generated from prior distributions of the parameters. If the estimation of the parameters is unbiased, posterior quantiles of the parameters obtained from PODs should be uniformly distributed (Cook et al., 2006; Wegmann et al., 2010). As with the empirical data, we calculated the posterior quantiles of true parameters for each pseudo run based on the posterior distribution of the regression-adjusted 1,000 simulations closest to each pseudo-observation.

**Supplemental TABLES**

**TABLE S1** Locality, code, number of genotyped individuals (*n*), latitude, longitude, elevation and genetic diversity (π, nucleotide diversity) for each studied population of *Omocestus bolivari* and *O. femoralis*.

| Species | Locality | Code | *n* | Latitude | Longitude | Elevation | π |
| --- | --- | --- | --- | --- | --- | --- | --- |
| *O. bolivari* | Sierra de La Pandera | PAND | 8 | 37.636927 | -3.805924 | 1510 | 0.0488 |
| *O. bolivari* | Sierra Mágina | MAGI | 8 | 37.740555 | -3.446550 | 1930 | 0.0573 |
| *O. bolivari* | Sierra de Arana | ARAN | 7 | 37.331304 | -3.516227 | 1736 | 0.0640 |
| *O. bolivari* | Albergue Universitario | UNIV | 6 | 37.096398 | -3.389056 | 2446 | 0.0726 |
| *O. bolivari* | Pozo de la Nieve | POZO | 6 | 37.368102 | -2.849309 | 2040 | 0.0739 |
| *O. bolivari* | Sierra de Baza | BAZA | 8 | 37.227338 | -2.752014 | 1876 | 0.0685 |
| *O. bolivari* | Sierra de Los Filabres | FILA | 7 | 37.219673 | -2.530702 | 2117 | 0.0637 |
| *O. bolivari* | Puerto de La Ragua | RAGU | 8 | 37.116477 | -3.026024 | 2100 | 0.0724 |
| *O. bolivari* | Loma de Piedra Blanca | LOMA | 8 | 36.975003 | -3.319273 | 2258 | 0.0718 |
| *O. bolivari* | Sierra de Lújar | LUJA | 8 | 36.836947 | -3.399291 | 1750 | 0.0621 |
| *O. bolivari* | Sierra de Gádor | GADO | 8 | 36.884610 | -2.803412 | 2077 | 0.0646 |
| *O. femoralis* | Pico Almenara | ALME | 7 | 38.543520 | -2.440236 | 1620 | 0.0796 |
| *O. femoralis* | Sierra de Cazorla | CAZO | 8 | 37.815379 | -2.959905 | 1814 | 0.0994 |
| *O. femoralis* | Poyotello | POYO | 8 | 38.119515 | -2.616541 | 1600 | 0.1050 |
| *O. femoralis* | Sierra de María | MARI | 5 | 37.676456 | -2.183102 | 1882 | 0.0884 |
| *O. femoralis* | Sierra de Espuña | ESPU | 8 | 37.865166 | -1.571250 | 1514 | 0.0639 |

**Supplemental figures**

**Figure S1** Root mean square error (RMSE) of parameter estimates against the number of partial least squares (PLS) components under six demographic models for (A) *Omocestus bolivari* and (B) *O. femoralis*.

**Figure S2** Number of reads per individual before and after different quality filtering steps by stacks for (A) *Omocestus bolivari* and (B) *O. femoralis*. The total height of the bars represents the total number of raw reads obtained for each individual. Within each bar, the dark red color represents the reads that were discarded by *process_radtags* due to low quality, adapter contamination or ambiguous barcode and orange color represents the reads that were discarded by *ustacks* after filtering out repetitive elements and reads that did not comply the different criteria required to create a “stack”. Green color represents the number of retained reads used to identify homologous loci. Population codes as in Table S1.

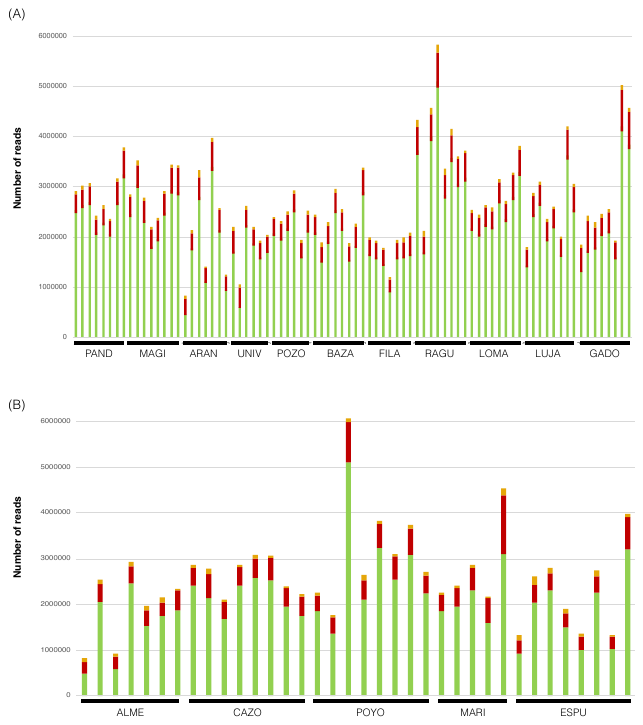

**Figure S3** Results of structure analyses for (A) *Omocestus bolivari* and (B) *O. femoralis*. The plots show mean (±SD) log probability of the data (Ln Pr(X|*K*)) over 10 runs of structure (left axes, solid dots and error bars) for each value of *K* and the magnitude of Δ*K* (right axes, open dots).

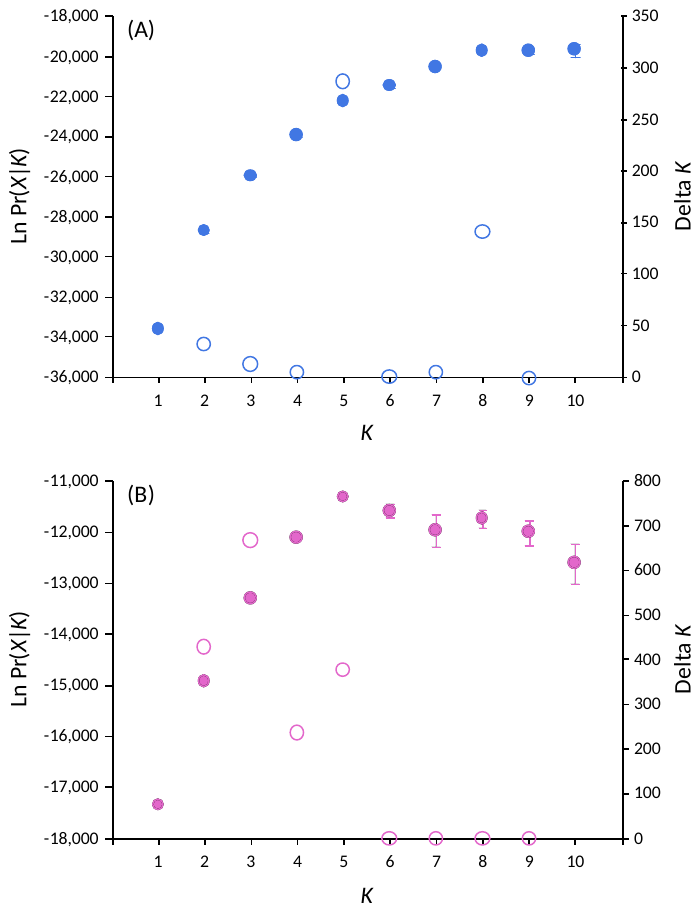

**Figure S4** Genetic assignment of individuals based on the results of structure for (A) *Omocestus bolivari* (from *K* = 2 to *K* = 8) and (B) *O. femoralis* (from *K* = 2 to *K* = 5). Individuals are partitioned into *K* colored segments representing the probability of belonging to the cluster with that color. Thin vertical black lines separate individuals from different populations. Population codes as in Table S1.

**Figure S5** Principal component analysis (PCAs) of genetic variation for (A) *Omocestus bolivari*  and (B) *O. femoralis*. Colors indicate the main genetic cluster at which individuals were assigned according to structure analyses for *K* = 8 (*O. bolivari*) and *K*= 5 (*O. femoralis*). Population codes as in Table S1.

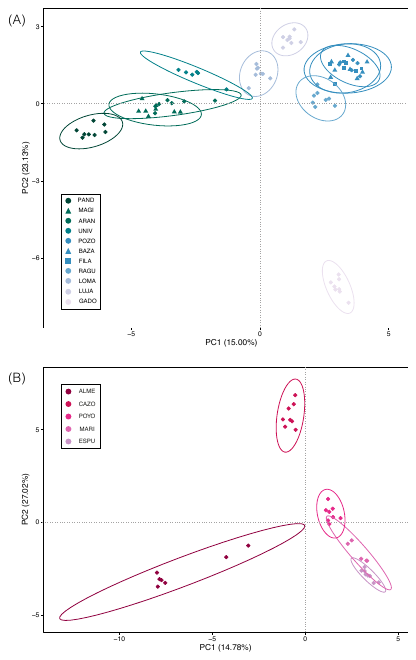

**FIGURE S6** Posterior distribution (solid black line) and mode (vertical dotted black line) of parameter estimates (*K*_MAX_, *m*, *N*_ANC_) for the most supported model for (A) *Omocestus bolivari* and (B) *O. femoralis* based on a general linear model (GLM) regression adjustment of the 1,000 retained simulations (0.5%) closest to empirical data. The comparison of posterior distributions before (blue shading) and after (solid black line) the ABC-GLM shows the improvement that this procedure had on parameter estimates. The comparison of prior (red shading) and posterior (solid black line and blue shading) distributions demostrates that the data contained information relevant to estimating the parameters. *K*_MAX_, carrying capacity of the deme with highest suitability (100%); *m*, migration rate per deme per generation; *N*_ANC_, ancestral population size.

**Figure S7** Distribution of posterior quantiles of true parameter values from 1,000 pseudo-observed data sets (PODs) used to assess bias in parameter estimation for the most supported model for (A) *Omocestus bolivari* and (B) *O. femoralis*. Posterior quantiles (grey bars) are compared to a uniform distribution (dashed red line) using a Kolmogorov–Smirnov test. Significant *p*-values indicate a deviation from a uniform distribution and potential bias in parameter estimation. *K*_MAX_, carrying capacity of the deme with highest suitability (100%); *m*, migration rate per deme per generation; *N*_ANC_, ancestral population size.

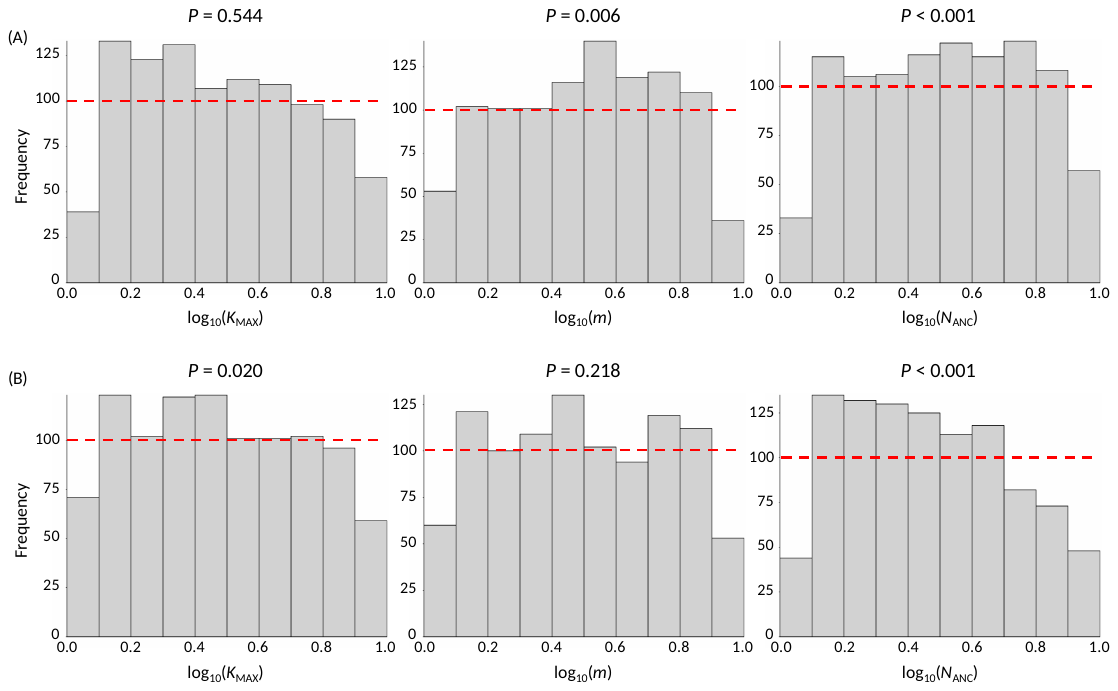
